# GSK-3α-BNIP3 axis promotes mitophagy in human cardiomyocytes under hypoxia

**DOI:** 10.1101/2024.05.24.595650

**Authors:** Hezlin Marzook, Anamika Gupta, Manju N. Jayakumar, Mohamed A. Saleh, Dhanendra Tomar, Rizwan Qaisar, Firdos Ahmad

**Author notes:** **Corresponding author: Firdos Ahmad, Ph.D.** Department of Basic Medical Sciences College of Medicine University of Sharjah Sharjah, UAE **Email;**.

## Abstract

Dysregulated autophagy/mitophagy is one of the major causes of cardiac injury in ischemic conditions. Glycogen synthase kinase-3alpha (GSK-3α) has been shown to play a crucial role in the pathophysiology of cardiac diseases. However, the precise role of GSK-3α in cardiac mitophagy remains unknown. Herein, we investigated the role of GSK-3α in cardiac mitophagy by employing AC16 human cardiomyocytes under the condition of acute hypoxia. We observed that the gain-of-GSK-3α function profoundly induced mitophagy in the AC16 cardiomyocytes post-hypoxia. Moreover, GSK-3α overexpression led to increased ROS generation and mitochondrial dysfunction in cardiomyocytes, accompanied by enhanced mitophagy displayed by increased mt-mKeima intensity under hypoxia. Mechanistically, we identified that GSK-3α promotes mitophagy through upregulation of BNIP3, caused by GSK-3α-mediated increase in expression of HIF-1α and FOXO3a in cardiomyocytes post-hypoxia. Moreover, GSK-3α displayed a physical interaction with BNIP3 and, inhibited PINK1 and Parkin recruitment to mitochondria was observed specifically under hypoxia. Taken together, we identified a novel mechanism of mitophagy in human cardiomyocytes. GSK-3α promotes mitochondrial dysfunction and regulates FOXO3a -mediated BNIP3 overexpression in cardiomyocytes to facilitate mitophagy following hypoxia. An interaction between GSK-3α and BNIP3 suggests a role of GSK-3α in BNIP3 recruitment to the mitochondrial membrane where it enhances mitophagy in stressed cardiomyocytes independent of the PINK1/Parkin.

## Introduction

Autophagy is an essential cellular process that helps in cellular turnover and delivering the cytoplasmic waste material to lysosomes for degradation. It is often considered as an adaptive mechanism that allows cells to survive under various stresses including nutrient deprivation and hypoxia. This process is executed through three main steps; 1) initiation of autophagosome formation, 2) expansion and formation of autophagosome complex, and finally 3) lysosomal fusion and degradation [1, 2]. Although conditions such as starvation and hypoxia induce modest autophagy, marked upregulation of lysosomal enzymes and enhanced autophagy occur post-myocardial infarction [3]. Mitophagy is autophagy of mitochondria in which autophagosomes selectively remove damaged or dysfunctional mitochondria [4]. This process is particularly important to maintain cardiac homeostasis as most of the energy required for the heart is generated by mitochondrial oxidative phosphorylation.

Glycogen synthase kinase-3 (GSK-3) has two closely similar isoforms, GSK-3α and GSK-3β, encoded by distinct genes [5]. Even though there is an overall 84% similarity between the two isoforms, they differ only by ∼2% within the catalytic domain [6, 7]. GSK-3α has a glycine-rich N-terminal region which distinguishes its biological function from the GSK-3β isoform [5]. Despite the large functional similarity between the two kinases, global loss of GSK-3α promotes cardiac hypertrophy, dysfunction and mortality post-myocardial infarction (MI), and aging [6–8]. Loss-of and gain-of-function in *in vivo* models have shown that GSK-3α plays a unique role in mitochondrial function and cardiac pathophysiology [9]. Accumulating evidences suggest that GSK-3α plays a crucial role in cardiac cell death in pathological conditions [9, 10]. Studies have shown that global genetic loss or pharmacological inhibition of GSK-3α leads to impaired autophagy, whereas blockade of the mTOR pathway restores autophagic markers in global GSK-3α knockout (KO) mice [11]. GSK-3α is also known to regulate ROS generation and mitochondrial function under stress conditions [12–15]. Hence, GSK-3α may play a role in cadiac mitophagy under pathological conditions.

BCL2/adenovirus E1B interacting protein 3 (BNIP3), an autophagy/mitophagy regulator, is also known to promote cardiac hypertrophy by mediating inflammatory response and signaling pathways responsible for abnormal mitophagy and apoptosis in cardiomyocytes. BNIP3 expression is modulated by several transcriptional factors and co-regulators such as FOXO3a under cardiac stress [16]. Studies have revealed that hypoxia-inducible factor 1α (HIF-1α) regulates BNIP3 gene expression during hypoxia. BNIP3 has LC3-binding motifs and translocates to the outer mitochondrial membrane, thereby recruiting autophagosomes to damaged mitochondria. FOXO3a-mediated activation of BNIP3 in cardiac oxidative stress induces an ER-mitochondrial Ca2+ shift, leading to mitochondrial and myocardial dysfunction [16, 17]. Thus, BNIP3 plays a constitutive role in removing dysfunctional mitochondria and investigating the role of GSK-3α in the regulation of these mitophagy pathways is important.

Here, we investigated the specific roles of GSK-3α in the regulation of mitophagy and related pathways in stressed cardiomyocytes. We show, for the first time, that GSK-3α promotes mitophagy in the human cardiomyocyte under acute hypoxia. GSK-3α induces BNIP3 expression through FOXO3a signaling mechanisms. Physical interaction of GSK-3α with BNIP3 likely facilitates its translocation to mitochondria. The translocated BNIP3 along with elevated LC3-II induces mitophagy in GSK-3α overexpressing cardiomyocytes post-hypoxia.

## Materials and Methods

### AC16 cardiomyocyte culture, transfection and treatments

AC16 cells were cultured under conditions previously described [18]. Briefly, the cells were grown in Dulbecco’s Modified Eagle’s Medium/Nutrient Mixture F12 (Sigma-Aldrich, #D6434) supplemented with 2 mM L-glutamine, 10% fetal bovine serum (FBS), and 1% penicillin/streptomycin antibiotic cocktail (Sigma-Aldrich, #P4333). The culture flasks were maintained in a humidified incubator at 37°C with 5% CO2 and 95% air. Cells were passaged at 80-90% confluence using 1X trypsin (Sigma-Aldrich, #T4299).

For transfection experiments, 0.32 x 10^6^ AC16 cells were seeded onto 6-well plates. Once cells reached approximately 60% confluence, they were serum-starved for 3 hours using Opti-MEM (Gibco, #00448). Subsequently, they were transfected with either control plasmid or Flag-GSK-3α WT human plasmid using Fugene 6 (Promega, #E2693) at a 3:1 ratio (DNA:Fugene 6). The transfection mixture was incubated with the cells for 3 hours in a CO2 incubator. After this period, an equal volume of complete DMEM containing double the usual concentration of FBS and penicillin-streptomycin was added to the cells, followed by further incubation for 24 hours in the CO2 incubator. To monitor LC3 conversion, pMRX-IP-GFP-LC3-RFP-LC3ΔG plasmid (Addgene plasmid # 84573) [19] was transfected according to the manufacturer’s instructions using Fugene 6. For studies investigating mitophagy, pCHAC-mt-mKeima vector (Addgene plasmid # 72342) [20] was transfected following the supplier’s protocol. Following the 24-hour transfection period, cells were subjected to the desired treatments.

Hypoxia was induced in AC16 cardiomyocytes using a regulated Whitley H45 hypoxic chamber and incubator. Before subjecting the cells to hypoxic conditions, the culture medium was replaced with serum-free media, and the cells were incubated for 3 hours in a CO2 incubator for acclimatization. Subsequently, cells were transferred to the hypoxic chamber with a controlled atmosphere of 95% N2, 5% CO2, and 1% O2, which was maintained throughout the hypoxic period. Unless otherwise specified, all experiments involved a 24-hour hypoxic incubation. To inhibit autophagy under the described conditions, cells were treated with 5 nM BafilomycinA1 (BafA1) (Sigma Aldrich, #B1793-10UG) for 24 hours.

### FACS analysis for GFP-LC-3-RFP conversion

Cells transfected with the plasmid were washed, trypsinized and collected in 200µL of 1X PBS. Cells were strained and subjected to FACS analysis (BD FACS ARIA III). Data acquisition was performed using forward scatter (FSC) for GFP and side scatter (SSC) for RFP components using linear scales and calibrated to exclude unstained cells and cellular debris. Mean fluorescence intensity (MFI) was measured using Flowjo software and graphs were plotted.

### Reactive Oxygen Species Assay

AC16 cells were seeded on 12-well tissue culture plates and transfected with GSK-3α plasmid. They were then subjected to hypoxia using the protocol described above. To assess cellular ROS generation, cells were harvested, washed, and resuspended in 100 μL of phosphate-buffered saline (PBS). The cells were then stained with 10 μM 2’,7’-Dichlorofluorescin diacetate (DCFDA) for 30 minutes. Fluorescence intensity was measured using a BD FACS Aria III flow cytometer, and the data was analyzed using FlowJo software. The median fluorescence intensity (MFI) was used to quantify ROS levels, and the results were presented graphically.

### Cell lysate preparation and immunoblotting

Following treatment, AC16 cardiomyocytes were kept on ice and washed twice with chilled PBS (Sigma-Aldrich, #8537). After removing the PBS from each culture plate, the cells were lysed by adding 1X RIPA lysis buffer (Thermo-scientific, #89900) supplemented with a phosphatase and protease inhibitor cocktail (Abcam, #ab201114) directly to the plate. Cells were then scraped to collect the lysate, which was transferred to microcentrifuge tubes. The lysate was centrifuged at 14,000 RPM for 20 minutes at 4°C. The supernatant was collected into a fresh tube, and protein concentration was determined using a colorimetric protein assay kit (BioRad, #5000006).

Western blotting was performed following previously described protocols [21]. Briefly, 20 μg of protein lysate was loaded onto an SDS-polyacrylamide gel and subjected to electrophoresis (BioRad). The proteins were then transferred from the gel onto a nitrocellulose membrane using a semi-dry method (BioRad Trans-Blot Turbo). The membrane was blocked with 5% non-fat dry milk in 1X TBST for 1 hour. This was followed by overnight incubation at 4°C with the primary antibodies listed in **Supplemental Table 1**. The next day, the membrane was washed three times with TBST solution for 5 minutes each. Subsequently, it was incubated with the corresponding HRP-conjugated secondary antibodies (Cell Signaling Technologies) for 1 hour on a shaker at room temperature. After further washes with TBST, the membrane was briefly incubated with Clarity Western ECL Substrate kit (Bio-Rad, #170-5060). Finally, the blots were imaged using a gel documentation system (BioRad Chemidoc Touch Imaging System). Band intensities were quantified using ImageJ software, and all samples were normalized to GAPDH levels.

### RNA extraction, cDNA preparation and qPCR

Total RNA was extracted from AC16 cells using the RNeasy kit (Qiagen, #74106) according to the manufacturer’s instructions. Briefly, cells were lysed directly with a buffer containing beta-mercaptoethanol, followed by washes with ethanol to purify the RNA. The isolated RNA was then eluted and dissolved in RNase-free water. The quality of the extracted RNA was assessed by measuring the absorbance ratios of OD260/OD280 and OD260/OD230 using a NanoDrop spectrophotometer. Subsequently, complementary DNA (cDNA) was synthesized from the RNA using the iScript cDNA Synthesis kit (BioRad, #170-8891). Quantitative real-time PCR (qRT-PCR) was performed using SYBR Green PCR master mix and primers specific for the genes of interest listed in the supplementary table (**Suppl. Table 2**). Gene expression levels were calculated using the delta-delta-Ct method and normalized to the housekeeping gene GAPDH.

### Immunofluorescence and Microscopy

AC16 cells were seeded on coverslips in 12 well-plates and transfected with plasmids. After respective treatments, cell were washed with PBS and fixed with 4% paraformaldehyde for 20 min at 37°C. Cells were then washed thrice with PBS. For immunostaining, cells were permeabilized with 0.01% TritonX for 10 min, followed by blocking with 3% BSA in 1XPBS with 0.01% TritonX. Cells were then washed with PBS for 5min/thrice and incubated with the indicated primary antibody **(Suppl. Table 1)** overnight. Cells were washed with PBS the next day and stained with secondary Alexa-fluor 488 antibody (Cell Signaling Technologies #4412S) for 1hr/RT and counterstained with DAPI. Cells were washed thrice with 1XPBS and coverslips were mounted on slides using Fluoromount (Abcam # ab104139). Slides were dried overnight and images were captured using a Nikon Eclipse. Image analysis was performed using NIS software. Quantification was done using ImageJ software to measure Mean fluorescence intensity.

### Co-Immunoprecipitation

Co-immunoprecipitation (Co-IP) assay was performed using a kit as per the manufacturer’s instructions (Thermo Scientific-26149). Briefly, cells were counted and approximately 3.5 x 10^4^ cells were seeded in multi-well plates. Cells were subjected to hypoxia and respective controls were maintained as indicated in the results. Post-treatment, cells were lysed using IP lysis buffer. Antibody immobilization to the column was carried out. Antibodies for immunoprecipitation were used at 1: 100 dilution. Lysates were pre-cleared and incubated with resin conjugated with respective antibodies. Post overnight incubation at 4°C, elution was carried out with elution buffer at pH 2.8. Samples were prepared for immunoblotting using 5X sample buffer containing 100 mM DTT to a final concentration of 1X. Samples were gently mixed and heated for 5min at 95^0^C. Once samples were cooled to room temperature, immunoblotting was carried out as described above and blots were probed with indicated antibodies **(Suppl. Table 1).**

### Mitochondrial extraction

Mitochondrial extraction was performed following the manufacturer’s protocol (Abcam ab110170) as described previously [15]. Briefly, a total of 1 x 10^6^ AC16 cells were seeded and subjected to hypoxia for 24h. Cells were resuspended in hypotonic buffer, Reagent A, to a protein concentration of 5mg/ml, and incubated on ice for 10 min, followed by homogenization. After centrifugation the supernatants were collected. The remaining pellet was then resuspended in Reagent B, centrifuged and supernatant collected. Next, both supernatants were combined, mixed vigorously and centrifuged. The mitochondrial pellet was resuspended in Reagent C, supplemented with protease and phosphatase inhibitors. For the cytosolic extract, acetone precipitation was carried out using ice-cold acetone. The pellet was air dried for 30 min and then subjected to lysis using RIPA buffer supplemented with protease and phosphatase inhibitors. Supernatants were collected and used as cytosolic fractions for western blotting with indicated antibodies **(Suppl. Table 1)**.

### Statistical analysis

Statistical analysis was performed to assess differences between the data groups. An unpaired t-test was used to compare two groups, while a one-way ANOVA was employed for comparisons between multiple groups with one independent variable. A two-way ANOVA was utilized when the experiment involved two independent variables. All statistical analyses were conducted using GraphPad Prism software (Graph Pad Prism Software Inc., San Diego, CA). Data are presented as mean ± standard error of the mean (SEM).

## Results

### GSK-3α induces autophagy under hypoxia

GSK-3α has been shown to regulate cardiomyocyte death under a variety of cardiac stressor situations [9, 22]. To better understand the underlying mechanism, we assessed the expression of autophagic markers in human AC16 cardiomyocytes post-hypoxia. The ratio of LC3-II /LC3-I is the most reliable marker to assess autophagy activation and related processes. We observed that the LC3-II/LC3-I ratio was comparable between GSK-3α overexpressing and control AC16 cells under normoxia however, the ratio significantly increased in GSK-3α overexpressing cardiomyocytes in comparison to control (**Fig. 1A, B**). In agreement, a significantly decrease in the p62/Beclin-1 ratio was seen only in GSK-3α-overexpressing cardiomyocytes under hypoxic but not normoxic conditions (**Fig 1C**). Consistently, the mRNA expression level of LC3 was significantly upregulated and the level of p62 was downregulated in GSK-3α overexpressing cardiomyocytes post-hypoxia (**Fig. 1D, E**). These observations suggest that GSK-3α promotes induction of autophagy in cardiomyocytes under hypoxia.

**Figure 1:**
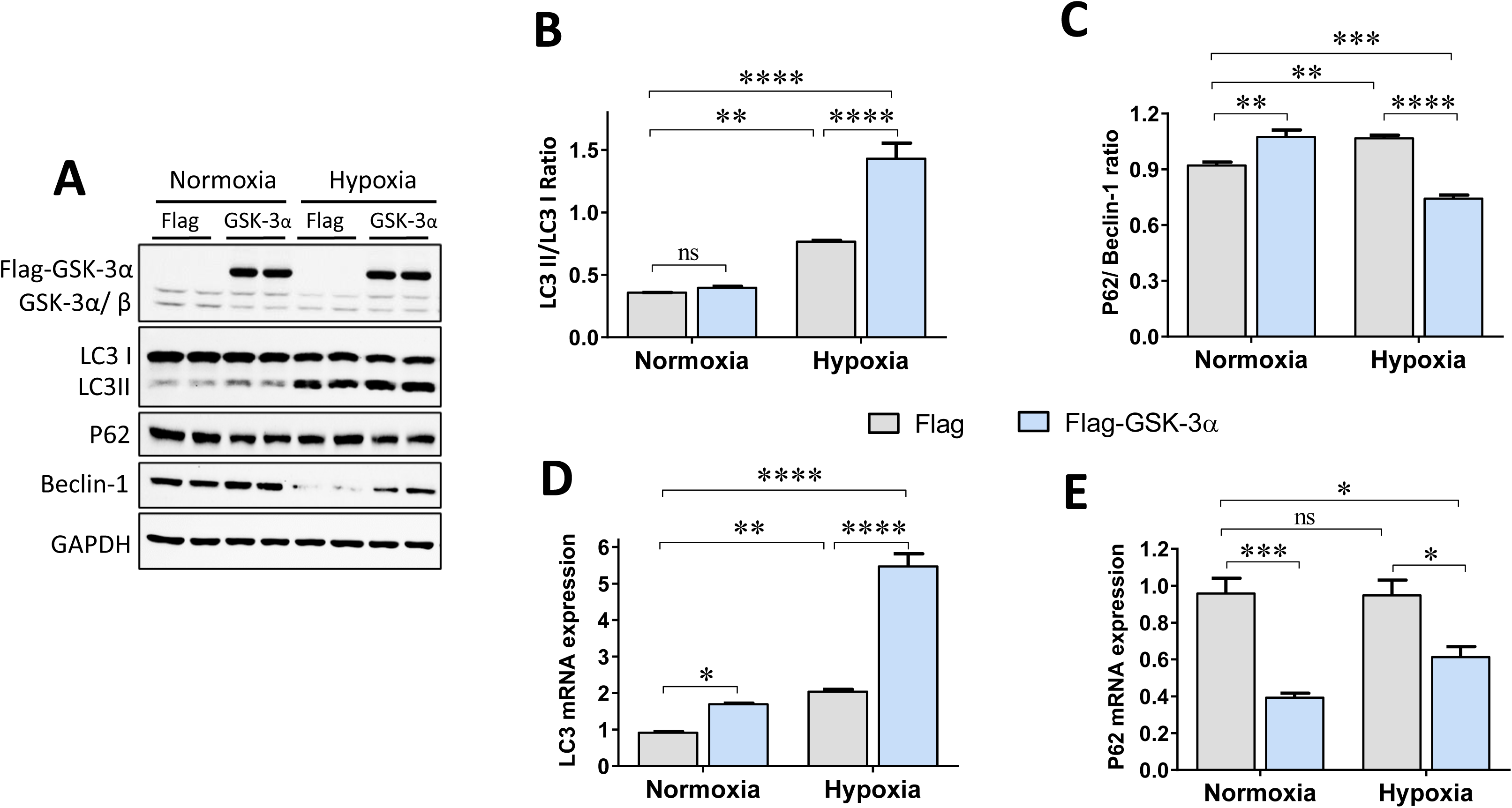
Autophagic flux in GSK-3α overexpressing cardiomyocytes post-24 hours of hypoxia: (**A**) Representative Western blots and quantification show (**B**) LC3 turnover and (**C**) Beclin-1 to p62 ratio. (**D-E**) Bar diagrams show mRNA expression levels of LC3 and p62 in GSK-3α overexpressing and control cells. *p<0.05, **p<0.01, ***p<0.001, ****p<0.0001.

### GSK-3α overexpression promotes autophagic flux in human cardiomyocytes under hypoxia

The increased LC3 cleavage and decreased p62/Beclin1 ratio in GSK-3α overexpressing cells post-hypoxia led us to examine LC3 turnover in more depth. LC3 overexpression was attained through transfection of GFP-LC3-RFP-LC3ΔG plasmid (**Suppl. Fig 1**) and autophagic flux was assessed, in which GFP-LC3 is degraded by autophagy, while RFP-LC3ΔG remains in the cytosol. The analysis of GFP and RFP intensity through flowcytometry showed a comparable GFP intensity in GSK-3α overexpressing cardiomyocytes compared to control under normoxia. The GFP intensity was significantly increased in hypoxia vs. normoxia group though the intensity was comparable between GSK-3α overexpressing cardiomyocytes and a Flag control when challenged with hypoxia (**Fig. 2A-B)**. In contrast, RFP intensity was comparable between control and overexpressing groups under normoxia. However, RFP intensity was significantly higher in GSK-3α overexpressing cardiomyocytes when compared to control under hypoxia (**Fig. 2 C**). These findings were further evaluated using confocal microscopy. Consistently, GFP intensity was found to be comparable between Flag and Flag-GSK-3a groups under both normoxia and hypoxia. However, the intensity of RFP was significantly higher in the GSK-3α overexpressing cells compared to the control cells post-hypoxia (**Fig 2D-F**). This suggests that under hypoxic conditions GSK-3α promotes autophagic flux and converts LC-GFP to RFP. These findings provide further evidence that the gain-of-GSK-3α function in human cardiomyocytes promotes autophagy under hypoxia.

**Figure 2:**
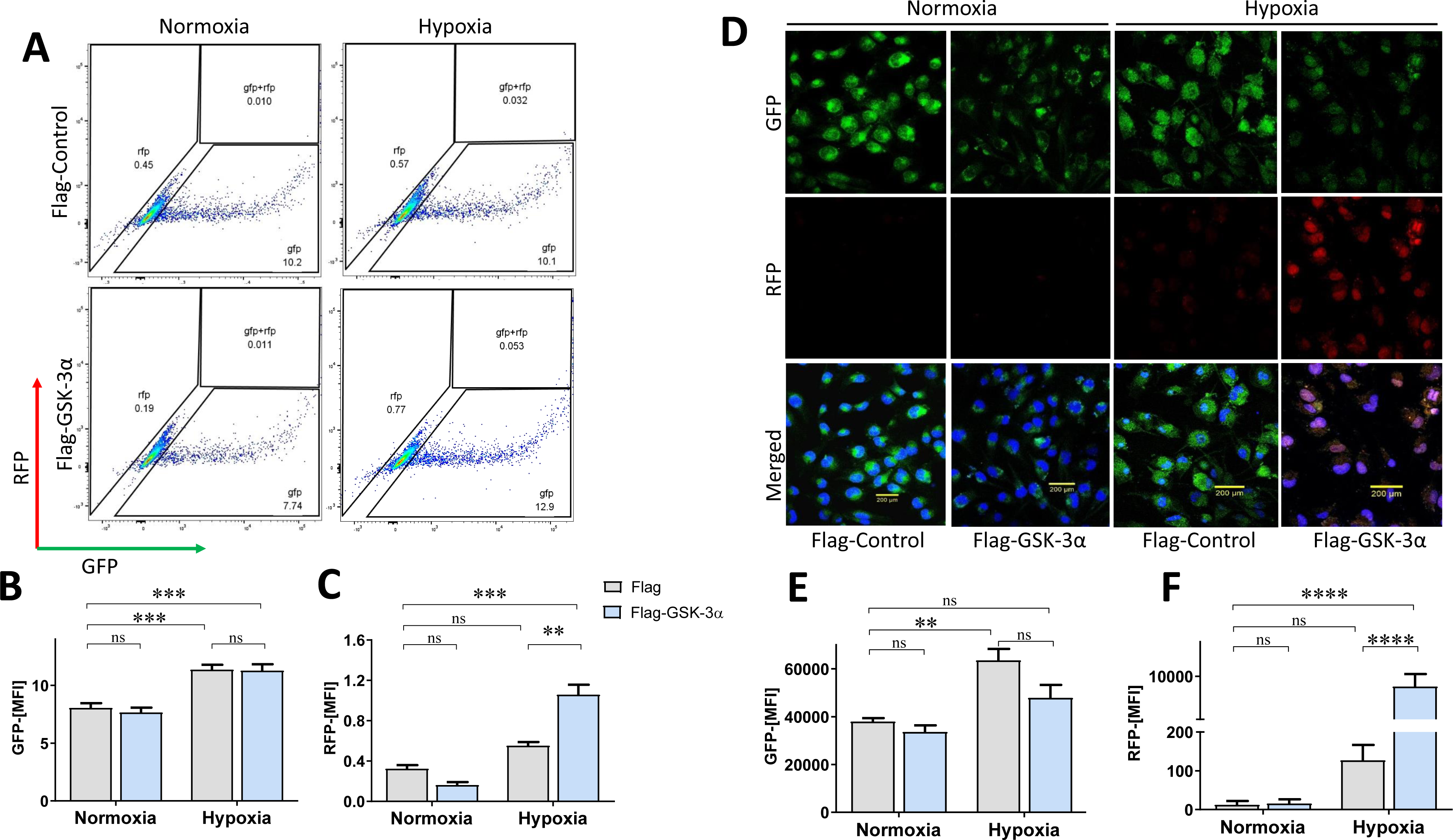
LC3 conversion in cardiomyocytes transfected with pMRX-IP-GFP-LC3-RFP-LC3ΔG plasmid: (**A**). Representative scatter plots show GFP, RFP and double-positive cell populations transfected with pMRX-IP-GFP-LC3-RFP-LC3ΔG plasmid. (**B-C**) Quantification shows the mean fluorescent intensity (MFI) of GFP and RFP. (**D**) Representative confocal microscopy images for pMRX-IP-GFP-LC3-RFP-LC3ΔG plasmid to detect LC3 conversion. Conversion of GFP to RFP signal represents an increased turnover of LC3-I to LC3-II. (**E-F**) Bar diagrams show the MFI of GFP and RFP. **p<0.01, ***p<0.001, ****p<0.0001.

To confirm the autophagic flux through LC3 cleavage post-hypoxia in detail, we treated GSK-3α overexpressing cardiomyocytes and control cells with autophagy inhibitor Bafilomycin A1 (BafA1), along with hypoxia and compared with the respective normoxic control. The effect of this treatment was also compared to the cells which underwent severe serum starvation (18h) induced autophagy under hypoxia. Autophagic marker analysis revealed profoundly inhibited LC3 cleavage and decreased p62 expression in GSK-3α overexpressing and control cells treated with BafA1 *vs.* untreated cells under normoxia. However, interestingly, BafA1 treatment largely inhibited LC3 cleavage and induced accumulation of p62 only in the GSK-3α overexpressing cardiomyocytes not in control cells post-hypoxia (**Fig 3A**). This is indicated by significantly reduced LC3-II/LC3-I ratio in GSK-3α overexpressing cardiomyocytes treated with BafA1 under hypoxia **(Fig 3B)**. Consistently, Beclin1 expression was reduced in GSK-3α overexpressing cells post-hypoxia *vs.* normoxia upon BafA1 treatment. The p62/Beclin1 ratio in GSK-3α overexpressing cells treated with BafA1 remained similar to control cells (**Fig. 3C**). These findings further support that GSK-3α promotes autophagic flux mainly through increased LC3 cleavage under hypoxia.

**Figure 3:**
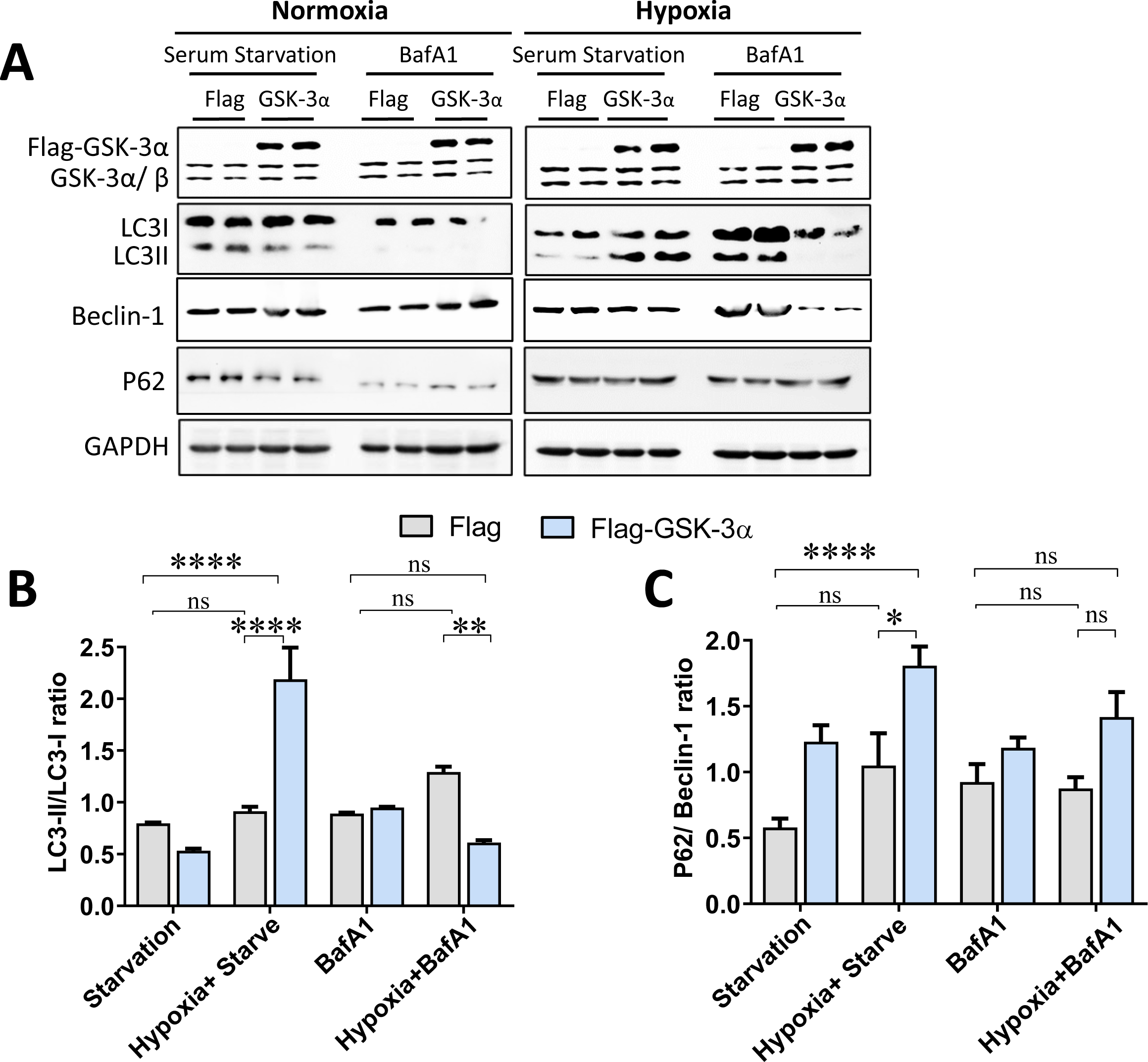
BafA1 treatment inhibits LC3 cleavage in GSK-3α overexpressing cardiomyocytes post-hypoxia: (**A**) Representative western blots for autophagy markers post-normoxia and hypoxia under prolonged starvation as well as BafilomycinA1 (BafA1) treatment. Bar diagrams show quantification for (**B**) LC-3II/LC3-I and (**C**) p62/Beclin-1 ratios. *p<0.05, **p<0.001, ****p<0.0001.

### GSK-3α promotes ROS generation and mitophagy

Since our results strongly suggest that GSK-3α promotes autophagy in a Beclin-1-independent manner and given the critical role of GSK-3α in the regulation of mitochondrial function in cardiomyocytes [8, 23], we hypothesized that mitophagy might play a role in this setting. In this series of experiments, we first evaluated the level of ROS generation in GSK-3α overexpressing cardiomyocytes post-hypoxia using DCFDA stain. The level of ROS was comparable between GSK-3α overexpressing *vs.* control cell group under normoxia. However, the level was significantly elevated in GSK-3α overexpressing cells compared to control cells under hypoxic conditions (**Fig. 4A-B**). Next, we assessed the role of mitochondria during hypoxia using the mt-mKeima vector (**Suppl. Fig. 2A-B**). In this model, cells experience acidic lysosome (pH 4.5) during which mt-Keima gradually shifts to longer-wavelength excitation under mitophagy. The AC16 cardiomyocytes were transfected with Flag or Flag-GSK-3α and mt-mKeima vectors and immunofluorescence staining was performed. The mean fluorescence intensity (MFI) showed a minimal intensity of mt-mKeima under normoxia however, a robust increased MFI was seen in the hypoxia *vs.* normoxia group irrespective of transfection type. The MFI was significantly higher in the Flag-GSK-3α overexpressing cells compared to Flag control cells under hypoxia (**Fig. 4C-D**). These results provide evidence that gain-of-GSK-3α function augments mitophagy in cardiomyocytes under hypoxia.

**Figure 4:**
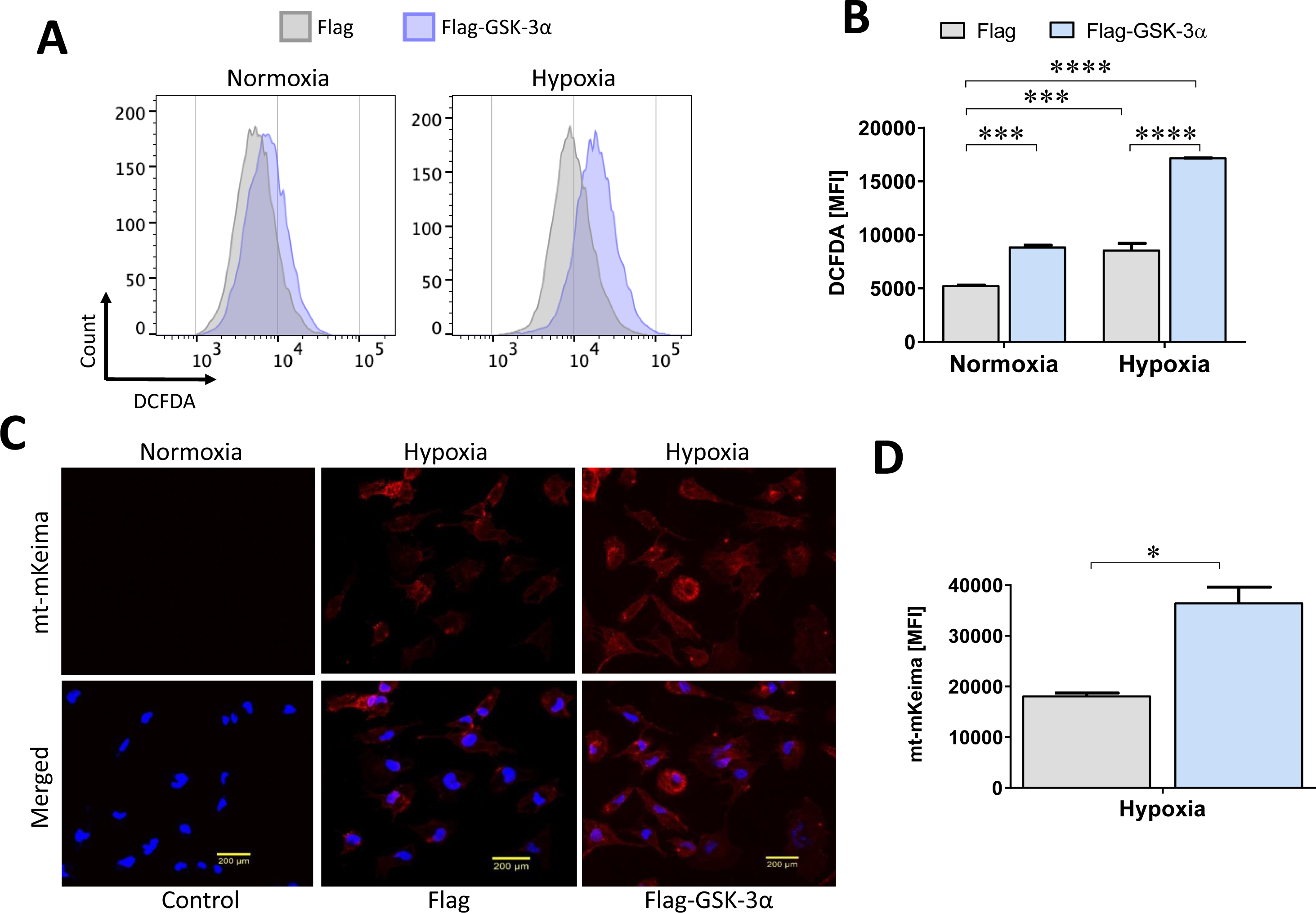
GSK-3α induces reactive oxygen species and mitochondrial autophagy in cardiomyocytes under hypoxia. (**A**) Representative flowcytometry histograms and (**B**) bar diagrams show the mean fluorescence intensity (MFI) of dichlorodihydrofluorescein diacetate (DCFDA) in control and GSK-3α overexpressing cardiomyocytes under normoxia and hypoxia measuring the levels of cellular reactive oxygen species (ROS). (**B**) Representative immunofluorescence images show the level of mt-mKeima in Flag and GSK-3α overexpressing cardiomyocytes under normoxia and hypoxia. (**C**). Quantification shows significantly higher MFI of mt-mKeima in GSK-3α overexpressing *vs.* Flag control cardiomyocytes. *p<0.05, ***p<0.001, ****p<0.0001.

### GSK-3α regulates mitophagy through BNIP3

Our findings indicate that GSK-3α regulates compensatory autophagy of dysfunctional mitochondria under hypoxia. Therefore, we sought to investigate the underlying mechanism and assess potential mitophagy regulatory partners. We identified BNIP3 as a key player in the GSK-3α -mediated mitophagy. BNIP3 expression at mRNA and protein levels was comparable in GSK-3α overexpressing compared to control cells under normoxia. However, the level of BNIP3 was profoundly elevated in the GSK-3α overexpressing cardiomyocytes vs. control cells challenged with hypoxia (**Fig. 5A, C, D**). Since FOXO3a is reported to regulate BNIP3 and mitochondrial function in the heart [16], we next assessed the potential involvement of FOXO3a. Indeed, the expression patterns of FOXO3a at both mRNA transcript and protein levels were similar to the pattern of BNIP3 in GSK-3α overexpressing cardiomyocytes under hypoxia **(Fig. 5B, C, E)**. Consistently, the level of HIF-1α, an upstream regulator of FOXO3a, was significantly upregulated in GSK-3α overexpressing cardiomyocytes post-hypoxia **(Fig. 5C, F)**.

**Figure 5:**
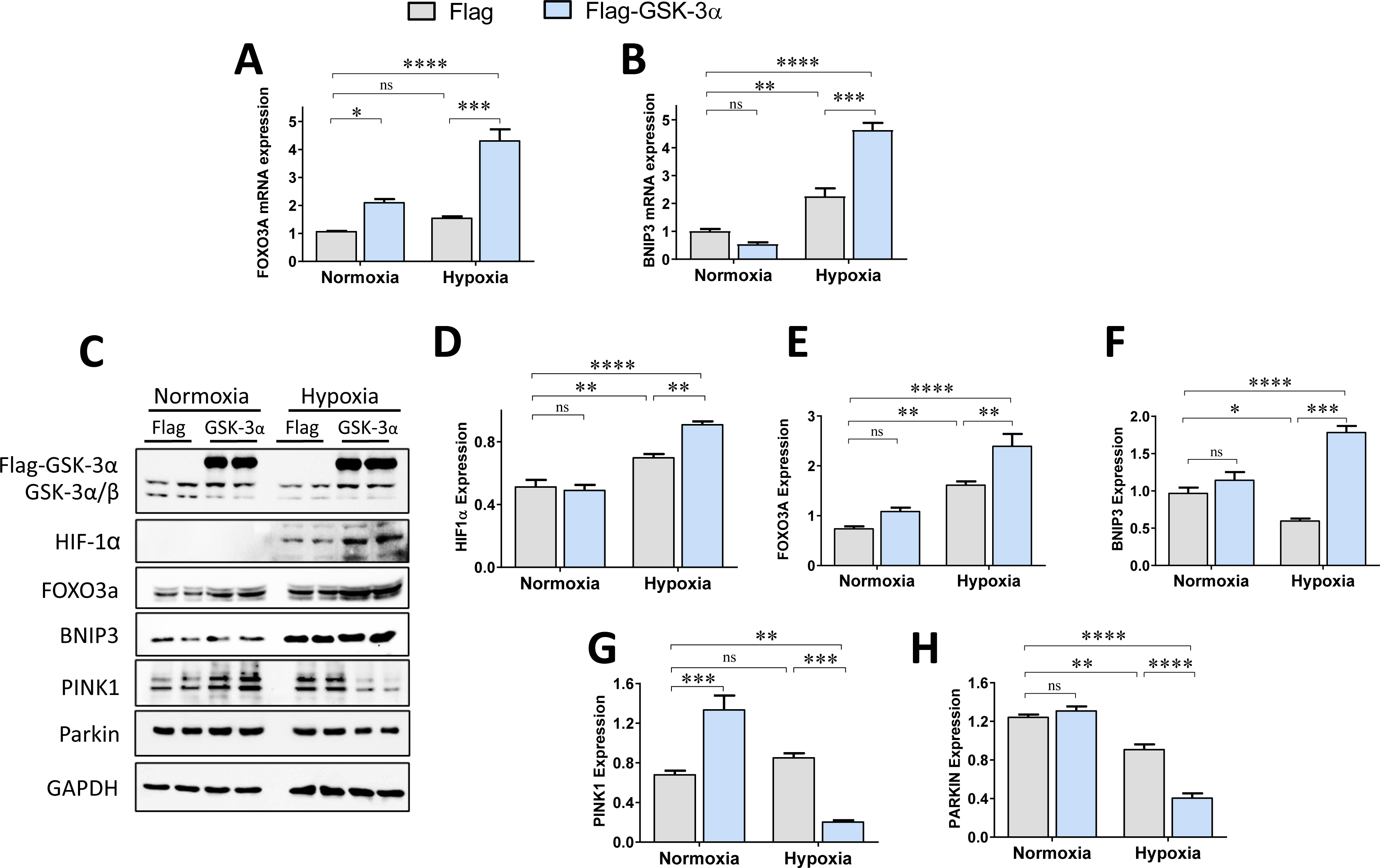
GSK-3α regulates BNIP3 in cardiomyocytes under hypoxia. (**A-B**) Bar diagrams show the relative mRNA expressions of BNIP3 and FOXO3a in Flag control and GSK-3α overexpressing cardiomyocytes under normoxia and hypoxia. (**C**). Representative Western blots and (**D-H**) quantification show the levels of GSK-3 isoforms, HIF-1α, FOXO3a, BNIP3, PINK1 and Parkin in control and GSK-3α overexpressing cardiomyocytes under normoxia and hypoxia. *p<0.05, **p<0.01, ***p<0.001, ****p<0.0001.

Recent evidence suggests a crucial role for Parkin and PTEN-induced kinase 1 (PINK1) in mitophagy [24] which is regulated by BNIP3 [25]. Therefore, we investigated the potential involvement of Parkin and PINK1 signaling in GSK-3α mediated mitophagy in cardiomyocytes. Gain-of-GSK-3α function induced the level of PINK1 and Parkin in cardiomyocytes under normoxia. However, in contrast and, inconsistent with FOXO3a/BNIP3 levels, the levels of both PINK1 and Parkin were downregulated in GSK-3α overexpressing compared to control cardiomyocytes post-hypoxia **(Fig. 5C, G, H)**. These findings suggest that GSK-3α-mediated mitophagy post-hypoxia is primarily regulated by BNIP3/FOXO3a signaling and PINK1/Parkin signaling plays a minimal role.

### GSK-3α physically interacts with BNIP3 and downregulates PINK1 and Parkin levels in mitochondria under hypoxia

To further investigate the precise role of GSK-3α in mitophagy, we isolated mitochondrial fractions from GSK-3α overexpressing cardiomyocytes. We observed the presence of GSK-3 isoforms both in mitochondrial as well as cytosolic fractions post-hypoxia (**Fig. 6A**). LC3 cleavage was significantly downregulated in mitochondria from gain-of-GSK-3α function samples under normoxia where the LC3-II/I ratio was significantly lower. In contrast, enhanced LC3 cleavage was observed in mitochondria from GSK-3α overexpressing cells under hypoxia (**Fig. 6B**). A similar pattern of LC3 cleavage was also observed in the cytoplasm of GSK-3α overexpressing cardiomyocytes challenged with hypoxia (**Fig. 6C**). The level of BNIP3 in the mitochondrial fraction was comparable in GSK-3α overexpressing cardiomyocytes *vs.* controls under normoxia; however, the level of BNIP3 was significantly upregulated in mitochondria from GSK-3α overexpressing cells post-hypoxia (**Fig. 6D**). Similarly, the BNIP3 level was also found to be elevated in the cytosol of GSK-3α overexpressing cells post-hypoxia (**Fig. 6E**). Consistent with BNIP3, the level of HIF-1α, an upstream FOXO3a regulator, was also elevated in the cytoplasmic and mitochondrial fractions of GSK-3α overexpressing cardiomyocytes post-hypoxia (**Fig. 6F-G**).

**Figure 6.**
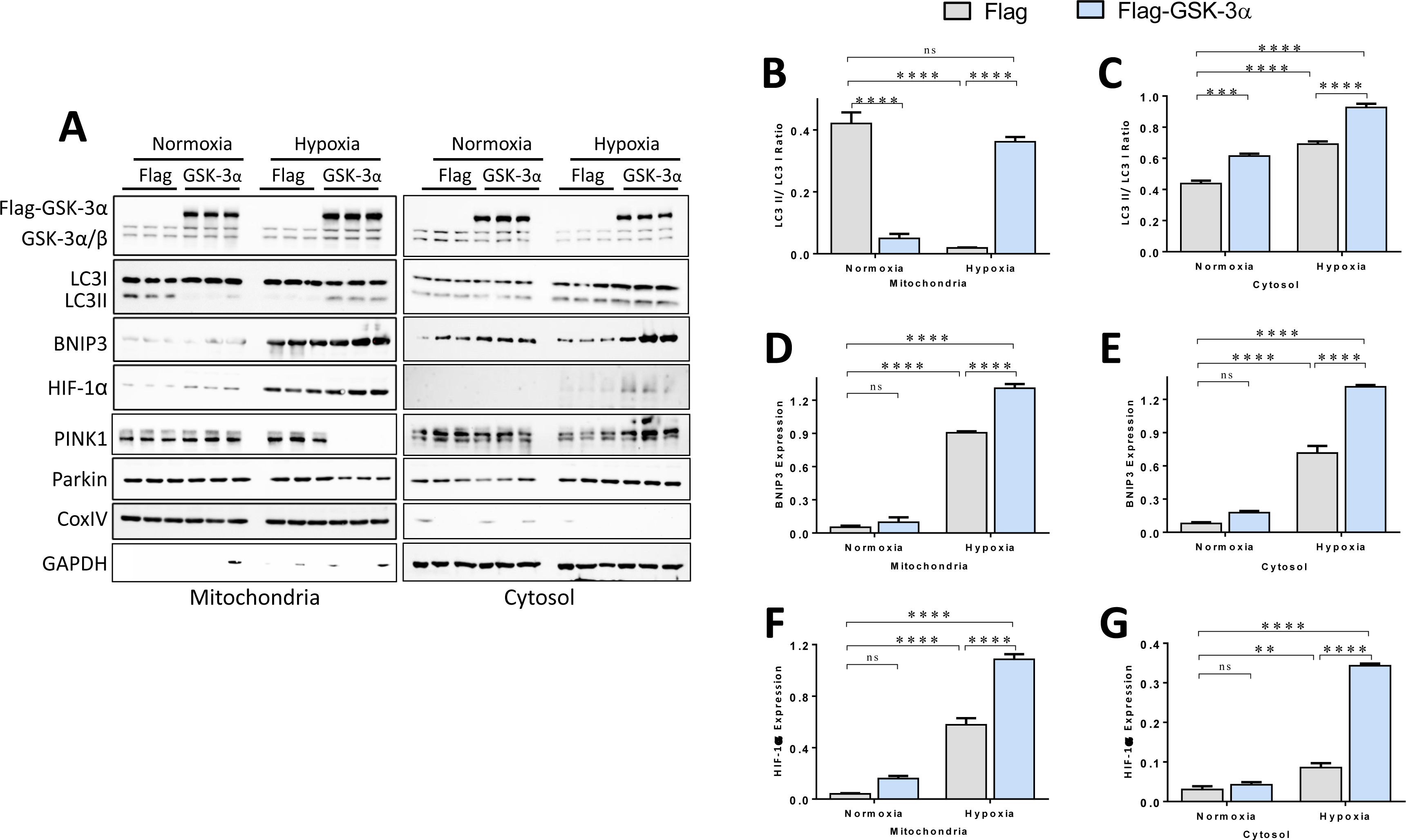
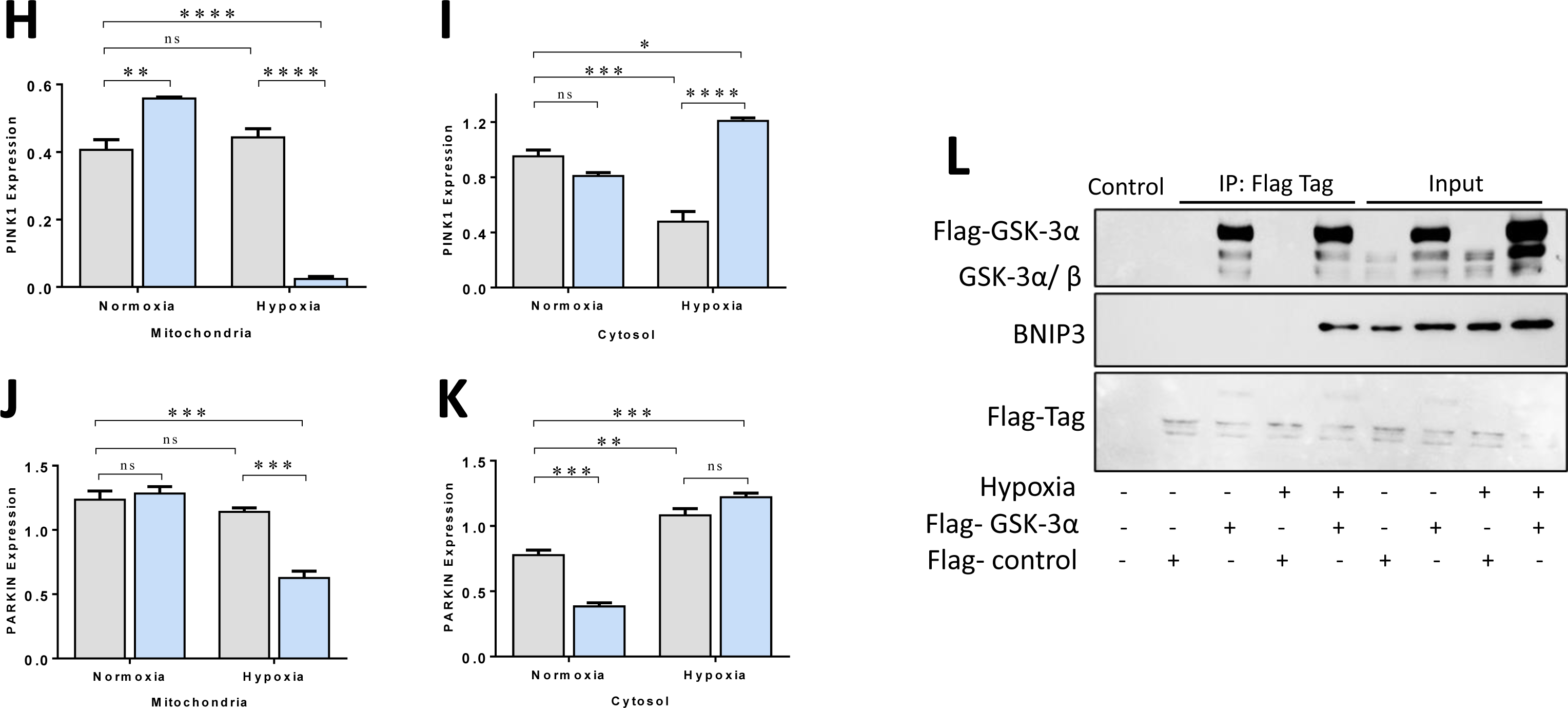
GSK-3α interacts with BNIP3 and regulates expressions of mitophagy markers under hypoxia. (**A**) Representative Western blots from mitochondrial and cytosolic (including nuclear) fractions of Flag and Flag-GSK-3α overexpressing cardiomyocytes, and (**B-K**) quantification show the relative expression of GSK-3 isoforms, LC3 turnover, BNIP3, HIF-1α, PINK1 and Parkin under normoxia and hypoxia. Blots from mitochondrial fraction were normalized by CoxIV and, blots from cytosolic fraction were normalized by GAPDH. (**L**) Co-immuno-precipitation blots show co-precipitation of BNIP3 with GSK-3α specifically under hypoxia condition when pulled down with Flag-tag antibody. *p<0.05, **p<0.01, ***p<0.001, ****p<0.0001.

Most strikingly, we observed translocation of PINK1 and Parkin between the cytoplasm and mitochondria particularly in GSK-3α overexpressing cardiomyocytes post-hypoxia. Lower levels of PINK1 and Parkin were observed in the mitochondria while the higher levels were detected in the cytoplasm of only GSK-3α overexpressing cells challenged with hypoxia but not in the normoxia group (**Fig. 6H-K**). These observations suggest that GSK-3α may limit PINK1 and Parkin recruitment to the mitochondrial membrane. To better understand this intriguing interplay between GSK-3α and BNIP3, we assessed whether GSK-3α physically interacts with BNIP3. Co-immunoprecipitation with Flag-tag antibody showed that GSK-3α associates with BNIP3 but only under hypoxic conditions (**Fig. 6L**). These observations lead us to conclude that GSK-3α plays crucial roles in the regulation of mitophagy in cardiomyocytes under hypoxia, possibly through physical interaction with BNIP3.

## Discussion

Mitochondrial dysfunction has been reported in a variety of cardiac pathological conditions and dysfunctional mitochondria can exert further detrimental effects on the heart. Mitophagy plays a crucial role in removing compromised mitochondria to prevent the generation of reactive oxygen species (ROS). Understanding the specific targets regulating mitophagy in the ischemic heart may offer potential strategies to enhance the removal of damaged mitochondria and promote mitochondrial renewal. Here, we have discovered that GSK-3α regulates cardiac mitophagy in a model of acute cardiac ischemia. GSK-3α overexpression in cardiomyocytes induces ROS generation and mitochondrial dysfunction in the cardiomyocytes post-hypoxia. Gain-of-GSK-3α function concomitantly promoted autophagic flux /mitophagy through increased LC3 cleavage and induction of Beclin1 expression, essential for autophagosome formation. Further, we identified a physical interaction between GSK-3α and BNIP3 which indicates regulation of mitophagy through BNIP3 **(Fig. 7)**. Moreover, GSK-3α -mediated translocation of PINK1/Parkin from mitochondria to the cytosol may further contribute to mitochondrial health in cardiomyocytes post-hypoxia.

**Figure 7:**
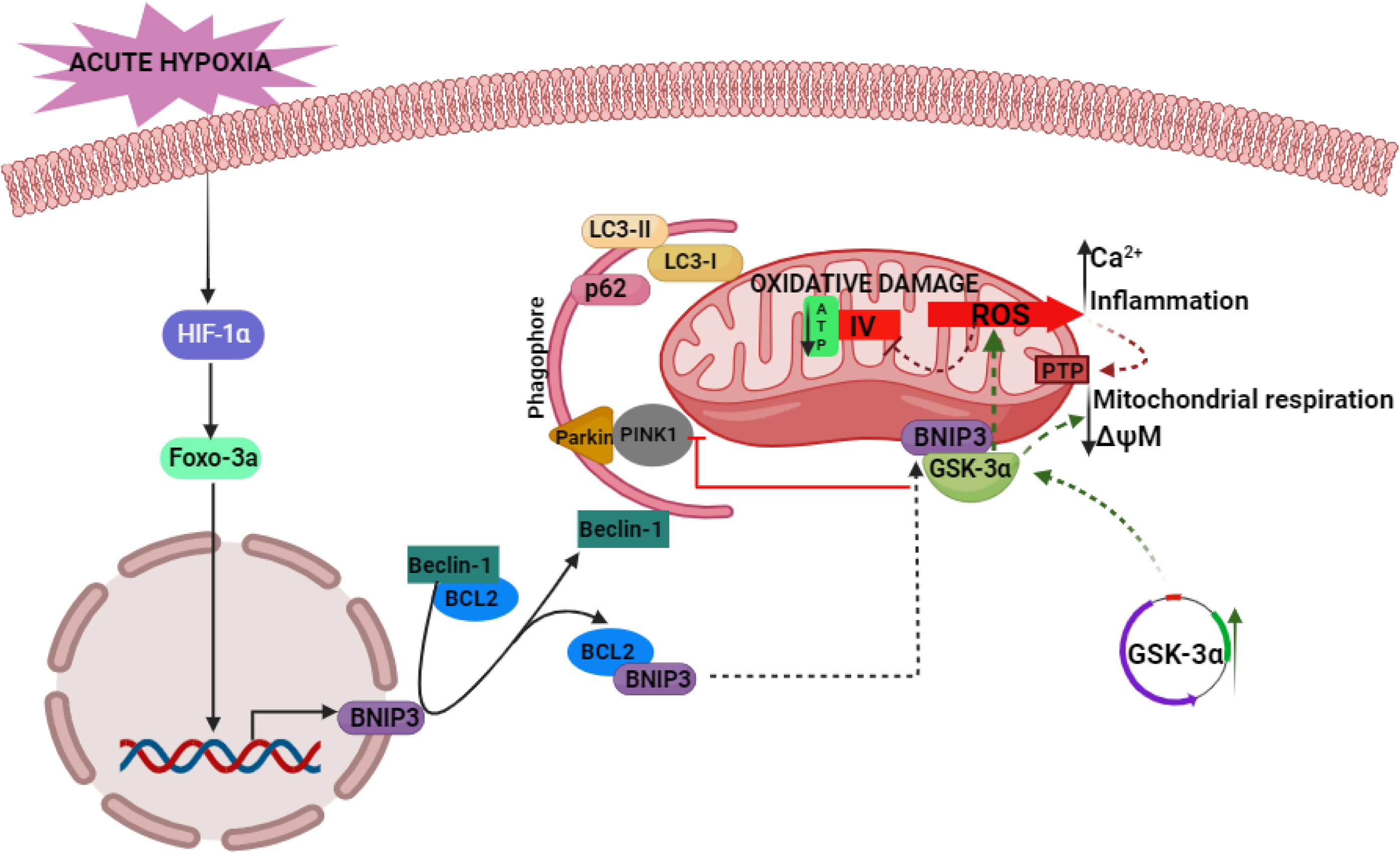
Schematic diagram depicts the role of GSK-3α in BNIP3-mediated mitophagy. Gain-of-GSK-3α function in cardiomyocytes induces levels of FOXO3a that ultimately upregulate BNIP3 expression under hypoxia. This induces the release of Beclin-1 from the Beclin-1-BCL2 complex. GSK-3α also induces LC3 cleavage in turn forming phagophore complex along with p62 and Beclin-1. GSK-3α physically interacts with BNIP3 and concomitantly inhibits PINK1 and Parkin recruitment to the mitochondrial membrane and causes PINK1-independent mitophagy in hypoxic cardiomyocytes.

Though various factors can contribute to mitochondrial damage in the heart under ischemic conditions, ROS is one of the major factors mediating cardiac tissue injury, followed by induction of mitophagy [26]. This cellular process further inhibits ROS generation by maintaining the quality control mechanism for detection and mitigation of mitochondrial dysfunction [27]. We observed elevated ROS generation in GSK-3α overexpressing cardiomyocytes post-hypoxia. Our recent report suggested that GSK-3α downregulates glutathione metabolism, possibly contributing to ROS generation and oxidative stress in cardiomyocytes post-hypoxia [15]. We observed an increased ROS generation associated with increased mitophagy in GSK-3α overexpressing cardiomyocytes post-hypoxia. These findings are consistent with a previous study that reported an interplay between ROS and mitophagy [28]. Mitophagy is not always protective, and its activation may facilitate cardiomyopathy [29]. We believe that the mitophagy activation in GSK-3α overexpressing cardiomyocytes post-hypoxia may be compensatory and activated to remove dysfunctional mitochondria from the dying cardiomyocytes.

Beclin-1 plays a key role in facilitating autophagy/mitophagy in the stressed heart [30]. Beclin-1, along with other required elements, senses elevated ROS levels and promotes autophagy to control excessive ROS accumulation and to remove dysfunctional mitochondria [31]. Consistent with these findings, we observed autophagy activation with marked increased LC3 cleavage and decreased p62 expression in GSK-3α overexpressing cells, particularly under hypoxia. The p62/Beclin-1 ratio is another valuable tool for monitoring autophagy/mitophagy activation [32] and a lower p62/Beclin-1 ratio in GSK-3α overexpressing cardiomyocytes under hypoxia further indicates activation of autophagy post-hypoxia. Interestingly, our study more specifically identified an increased mitophagy in GSK-3α overexpressing cardiomyocytes post-hypoxia which was confirmed by elevated mt-mKeima intensity, a marker of mitophagy. Consistent with these findings, excessive mitophagy was previously reported to cause myocardial damage [29]. Similarly, autophagy induction has also been reported to aggravate cardiac injury and dysfunction in mice with type-1 diabetes mellitus [33]. Akt, an upstream regulator of GSK-3α, plays a crucial role in the autophagy/ mitophagy. Akt2 deficiency induces mitophagy and promotes cell death by excessive removal of mitochondria [34]. In agreement with this, another study has demonstrated that inhibition of Akt signaling attenuates mitophagy by limiting PINK1 accumulation [35]. Akt signaling inhibition leads to downstream activation of GSK-3α and, consistent with our findings, observed increased mitophagy in these studies was potentially regulated through increased GSK-3α activity.

BNIP3 is another signaling molecule tightly regulated under normal physiological conditions but upregulated in response to various cellular stresses including oxidative stress caused by elevated ROS generation under hypoxia, and ischemia-reperfusion (I/R) [36, 37]. The upregulation and activation of BNIP3, along with other autophagy-related proteins, leads to the formation of autophagosomes around damaged mitochondria and induces mitophagy [38]. Consistent with this, we observed elevated levels of BNIP3 and LC3 cleavage in GSK-3α overexpressing cardiomyocytes and the mitochondrial fraction, compared to control cells, particularly under hypoxia but not normoxia. Moreover, a physical interaction between GSK-3α and BNIP3 supports a role for GSK-3α in BNIP3-mediated autophagosome formation and mitophagy in sick cardiomyocytes following hypoxia. Similar to our observations in hypoxia, a study has reported that BNIP3 promotes I/R-induced cardiac injury and consequently activates a protective stress response with the induction of mitophagy to remove damaged mitochondria [39]. Several other studies have demonstrated a detrimental role for BNIP3. Overexpression of BNIP3 causes loss of mitochondrial membrane potential, cytochrome-c release and, ultimately, mitochondrial dysfunction [40]. Another study reported that BNIP3 overexpression promotes mitophagy and cardiomyocyte apoptosis and that loss of BNIP3 reverses pressure overload-induced cardiac remodeling and diastolic dysfunction [41]. We observed an elevated level of ROS in GSK-3α overexpressing cardiomyocytes and increased ROS generation has been previously linked to HIF-1α-mediated BNIP3 induction and mitophagy activation [42]. HIF-1α regulates BNIP3 not only through direct binding with hypoxia-response elements within the BNIP3 promoter region but also through upregulation of FOXO3a [43]. Similarly, we observed an increased level of FOXO3a in GSK-3α overexpressing cardiomyocytes post-hypoxia. Thus, GSK-3α plays a crucial role in mitophagy activation through BNIP3 upregulation under hypoxia.

Cellular stress triggers mitophagy through multiple mechanisms. It can be either regulated through ubiquitin-dependent or -independent (or receptor-mediated) pathways [44]. In the ubiquitin-mediated pathway, generally referred as the PINK1/Parkin pathway, of mitophagy, PINK1 is stabilized on the outer mitochondrial membrane (OMM) following stress, which then promotes Parkin recruitment. Accumulating evidence suggests that excessive ROS can also activate the PINK1/Parkin signaling pathway to promote clearance of impaired mitochondria from cardiomyocytes [24]. An interaction between BNIP3 and PINK1 facilitates Parkin recruitment on the OMM and regulates mitophagy of compromised mitochondria under hypoxia [25, 45]. We did not observe PINK1 and Parkin recruitment onto mitochondria. Rather, lower levels of PINK1 and Parkin were observed in mitochondria compared to the cytosolic fraction of GSK-3α overexpressing cardiomyocytes post-hypoxia. This indicates for the regulation of receptor-mediated alternate pathway where BNIP3, associated with the OMM, can directly bind to LC3 and direct damaged mitochondria to autophagosomes for degradation and removal [4]. These findings strongly suggest that GSK-3α regulates BNIP3-mediated mitophagy with minimal or no involvement of PINK1/Parkin signaling.

Our findings align with a growing body of research highlighting the importance of protein kinases in PINK1/Parkin-independent mitophagy [46]. Beyond BNIP3, other protein such as Beclin1-regulated autophagy protein 1 (AMBRA1) has been shown to regulate PINK1/Parkin-independent mitophagy through the HUWE1 E3 ligase/IKKα pathway [47]. Additionally, recent studies suggest that lipid-binding kinases, cyclin G-associated kinase (GAK) and protein kinase C Delta (PRKCD), play critical roles in mitophagy independent of Parkin [48]. OMM proteins Nix and FUNDC1, which share structural similarities with BNIP3, have also been implicated in PINK1/Parkin-independent mitophagy [49, 50]. Furthermore, Drp1, a dynamin related GTPase, has been shown to mediate the selective elimination of damaged mitochondria independent of PINK1/Parkin. Drp1 directly interacts with ubiquitin chains on the OMM, promoting the removal of damaged portions, which are then engulfed by autophagosomes for degradation [51]. These findings are further supported by *in vivo* studies demonstrating that PINK1 is not essential for Parkin recruitment to mitochondria or for mitophagy activation in cardiomyocytes [52]. This further strengthens the evidence for PINK1-independent forms of mitophagy in cardiomyocytes, and GSK-3α might play a pivotal role in this alternative pathway.

In summary, our study reveals a novel role for GSK-3α in mitophagy induction in stressed human cardiomyocytes. Gain-of-GSK-3α function profoundly induces ROS generation and autophagic flux through increased LC3 cleavage and Beclin-1 expression in cardiomyocytes challenged with acute hypoxia. GSK-3α physically interacts with BNIP3 and, GSK-3α overexpression promotes mitophagy through activation of the FOXO3a/BNIP3/LC3 pathway in cardiomyocytes under hypoxia.

## Declaration

### Ethical Approval

N/A, no animals or human subjects were involved.

### Authors’ contributions

**HM** performed experiments, collected data, performed analysis and helped with manuscript writing, **AA** and **MNJ** performed experiments and collected data, **DT** helped with reagents, data analysis and manuscript editing, **MAS** and **RQ** helped with data analysis, interpretation and manuscript editing, **FA** designed the study, acquired funding, supervised the project, performed data analysis and interpretation, and wrote and revise the manuscript.

### Availability of Data

Data is included in the manuscript and supplementary files. Additional information is available from the corresponding author upon a reasonable request.

