## Supplementary file for "GSK-3α-BNIP3 axis promotes mitophagy in human cardiomyocytes under hypoxia"

**Supplementary Fig. 1**

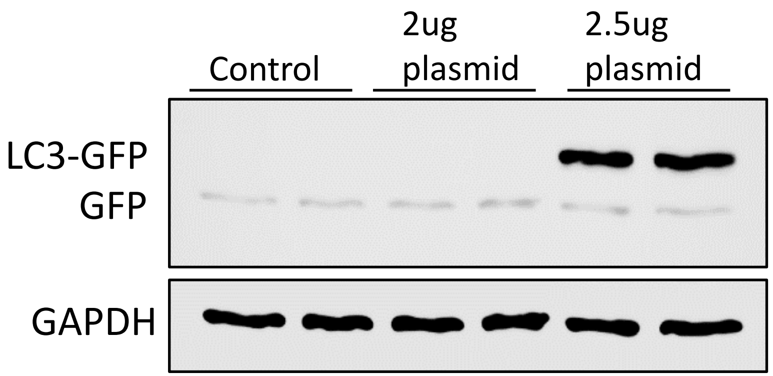


**Supplementary Fig 1:** standardization of pMRX-IP-GFP-LC3-RFP-LC3ΔG plasmid concentration in AC16 cardiomyocytes.

Supplementary Fig. 2

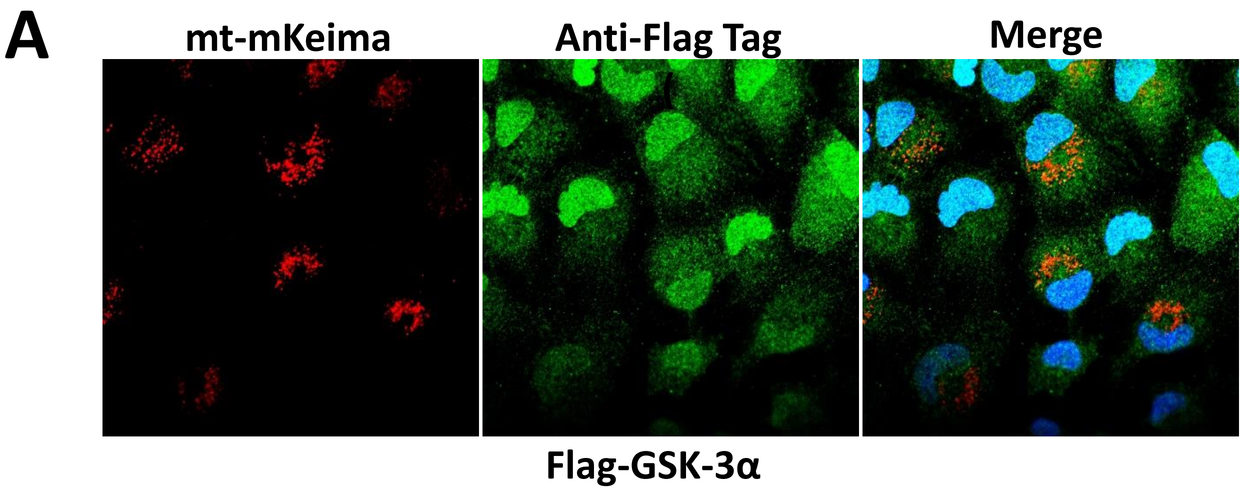


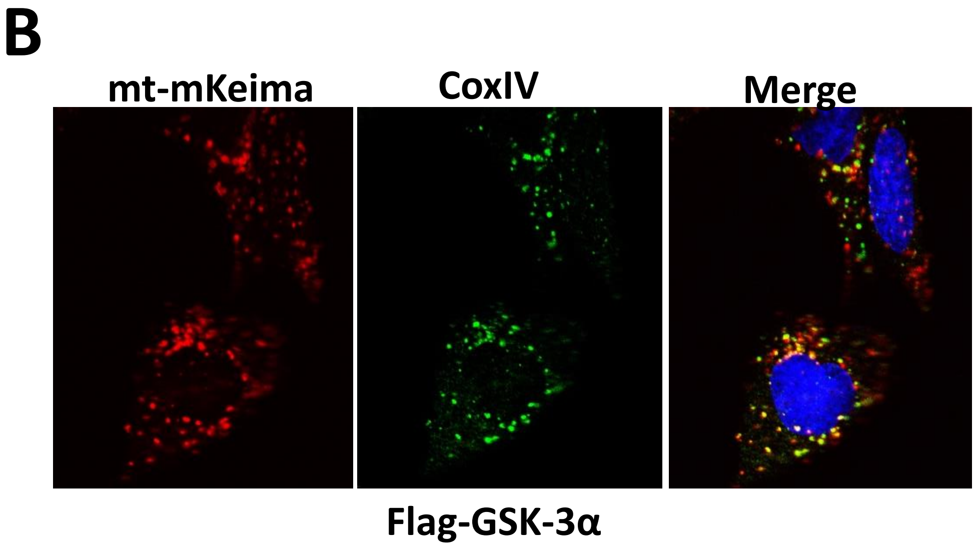


**Supplementary Fig 2:** (**A**) Validation of mt-mKeima plasmid to confirm the co-localization with CoxIV mitochondrial marker. (**B**) Immunofluorescence staining of Flag-tag antibody in GSK-3α overexpressing cardiomyocytes co-transfected with mt-mKeima. Red represents mitochondrial staining of mt-mKeima (mitophagy) under hypoxia and green represents GSK-3α.

**Supplementary table 1: List of antibodies:**

| | | |
|---|---|---|
| 1 | Anti-Flag Tag | Cell signaling technology #14793 |
| 2 | BECLIN1 | Cell Signaling Technology #3495 |
| 3 | BNIP3 | Cell Signaling Technology #3769 |
| 4 | COX-IV | Cell signaling technology #4850T |
| 5 | FOXO3a | Cell Signaling Technology #2497 |
| 6 | GSK-3α/β | Cell Signaling Technology #5676 |
| 7 | GAPDH | Fitzgerald #10R-G109A |
| 8 | GFP | Cell signaling technology #2955 |
| 9 | HIF-1α | Cell Signaling Technology #14179 |
| 10 | LC3 A/B | Cell Signaling Technology #12741 |
| 11 | PARKIN | Cell signaling technology #32833 |
| 12 | PINK1 | Cell signaling technology #6946 |
| 13 | SQSTM/P62 | Cell Signaling Technology #5114 |

**Supplementary table 2: List of primers:**

1. **FOXO3A**

Forward Sequence TCTACGAGTGGATGGTGCGTTG

Reverse Sequence CTCTTGCCAGTTCCCTCATTCTG

2. **BNIP3**

Forward Sequence TCAGCATGAGGAACACGAGCGT

Reverse Sequence GAGGTTGTCAGACGCCTTCCAA

3. **LC3B**

Forward Sequence GAGAAGCAGCTTCCTGTTCTGG

Reverse Sequence GTGTCCGTTCACCAACAGGAAG

4. **GSK3A**

Forward Sequence GCAGATCATGCGTAAGCTGGAC

Reverse Sequence GGTACACTGTCTCGGGCACATA
